# If Research is Not the Evidence, What is it? Egyptian Physicians’ Explanations of the Lack of Research Citations in their Health Vlogs

**DOI:** 10.1101/2022.04.25.489430

**Authors:** Noha Atef

## Abstract

Doctor-influencers have the knowledge needed to understand scientific research, and they have the social media popularity to share it with large audiences. In this article, I explore the treatment of evidence in vlogs by Egyptian doctor-influencers: how they present it and what they use instead of it. I answer the following questions: 1) How do doctor-influencers present evidence on social media? 2) If evidence is not the research, then what is it? The data was collected through in-depth interviews with 12 Egyptian doctor-influencers from different specializations; two focus groups; and a critical discourse analysis of 48 of the influencers’ most popular and most engaging videos. I found that the doctor-influencers cited academic research only in the videos about medical controversies, new treatments, misinformation, or common medical mistakes. In most of their videos, the Egyptian doctor-influencers scarcely referred to medical research. This is because they believe that their audiences would not understand evidence due to their low educational attainment, weak research skills, and the language barrier (as most medical research is published in English rather than Arabic). These findings highlight the challenges of communicating science in societies with a low level of education. In these settings, communicators may need to move beyond mediating research studies on social media and instead focus on simplifying, translating, and connecting evidence-based information to the cultural context.

## Introduction

While patient-centered communication (PCC) has long been endorsed as a key element of quality health care (Jiang & Liu, 2020), it is inaccessible to many people in the Global South. The shortage of physicians is one reason why PCC is hard to achieve in developing countries. For instance, in 2020, Egypt had 10.8 physicians per 10,000 inhabitants (World Health Organization, Global Health Statistics, 2020). This means that physicians receive many patients every day, which limits their communication with them (Metwally et al., 2021). In this case, seeking health information on the internet can improve the patient-physician relationship by making the patients “internet-informed” about their conditions (Langford et al., 2020). This helps patients exchange medical information with health-care professionals (Huo et al., 2019).

Many studies have identified social media as a source of health misinformation (see Suarez-Lledo & Alvarez-Galvez, 2021; Sylvia Chou et al., 2020; Vraga & Bode, 2020). There is also a growing body of research on doctor-influencers’ efforts to spread correct health information online; however, there is almost no scholarship on the use of evidence in different cultural contexts. This article explores the use of evidence in vlogging by Egyptian doctor-influencers, including its presentation and substitutes. I answer the following questions: 1) How do doctor-influencers present evidence on social media? 2) If evidence is not the research, what is it? In this article, “evidence” denotes the use of academic scholarship as the scientific foundation of health information, and “doctor-influencers” are the certified health professionals who are popular video bloggers (vloggers). It is essential to note this article does not evaluate the accuracy of information presented by the vloggers; it only investigates how they present this information.

The article focuses specifically on Egyptian doctor vloggers. Very little research on health communication is conducted in the Global South. Egypt is a unique case for studying online health communication because of the high mobile penetration in this country, which has reached 93.6% by December 2020, enabling 52 million people to use the internet through mobile devices (ICT Indicators Bulletin, Egyptian Ministry of Communication and Information Technology, 2020). On the other hand, the high illiteracy rate among Egyptians, reaching 29% (World Bank, 2017a), suggests low health literacy rate. A survey of 805 attendees of one university hospital in Cairo showed that 84% of them had limited functional health literacy—the ability to understand and act on medical instructions (Almaleh et al., 2017). At the same time, Egypt suffers from a notable shortage of medical capacity; in 2015, there were 1.5 beds per 1,000 people, and the number went down to 1.4 beds in 2017. In both cases, the figure is significantly below the world average of 2.7 beds per 1,000 people (World Bank, 2017b). Inequities persist based on income and across governorates (Shawki, 2018).

## Literature Review

### Doctor-Patient Communication

Szasz and Hollender (1956) identified three types of doctor-patient relationships based on the amount of information exchanged between the two parties according to the model of communication. The first is the active-passive model, where the doctor has all the authority to choose a treatment and execute it, while the patient is not informed about the rationale of treatment. The second is the guidance-cooperation model, where the physician decides on the remedy and informs the patient about it. The last one, mutual participation, is a patient-centered relationship. These models were further refined and elaborated by Emanuel and Emanuel (1992), who identified the following. 1) The paternalistic model: the physician determines the right treatment and gives the patient only such information as would encourage the patient to consent to the treatment, or just tells the patient when the treatment would be carried out. In this model, the physician has complete authority over the patient. 2) The informative model: the physician provides the patient with all information relevant to their condition, and the patient selects the medical interventions they want. Here, the physician informs the patient about all the risks and side effects of every treatment, and the patient makes the decision; therefore, this model is also called the consumer model. 3) The interpretive model: s similar to the informative model; however, in this model, the physician helps them with that. Thus, the physician is like a consultant for the patient, telling the patient about all the available treatments and providing the relevant information, as well as helping them select what they want. 4) The deliberative model: the physician acts as a friend or teacher to the patient as the two parties discuss the best treatment options.

The doctor-patient relationship is influenced by the economic situation; in developing countries with poor health services, the paternalistic model is dominant, and the patient-centered perspectives are likely to be minimally represented in the medical curriculum (Unger et al., 2003). People with high income and education have the highest chance of receiving patient-centered care (Hassig, 2021). Effectively, the physician-centered models are dominant in low- and middle-income countries. For instance, in China, the paternalistic model is common in doctor-patient communications (Jiang & Liu, 2020), where the patients play a passive role and avoid posing questions to their doctors (Jiang & Street, 2019). “Medical paternalism” is also observed in Sub-Saharan African countries (Norman, 2015); such as Nigeria where the culture grants the healthcare providers power over patients, which reflects in the physician-patient interactions (Adebayo, 2021). The paternalistic model also appears in Iran and was represented in the first and the only medical drama produced and broadcasted in Iran on the National TV Channel One with the sponsorship of the Ministry of Health and Medical Education (Riahi et al., 2020)

### Egyptian Doctor-Patient Communication

In Egypt, the discipline of doctor-patient communication generally does not receive enough attention. Shalaby et al. (2019) found that for Egyptian physicians, the patient’s satisfaction is the main indicator for the quality of their communication with the doctor; 49.5% of 275 randomly selected physicians agreed that “Effective doctor-patient communication is highly associated with increased patient satisfaction.” In the same study, 41.8% of participant physicians said they thought they had “good” communication with their patients, with correlation between the physicians’ years of experience and perceived communication skills. Generally, the rate of dissatisfaction with the physician-patient relationship is high among medical students; 75.9% of medical students were not satisfied with their relationship with patients (Kabbash et al., 2021). A relevant nuance is that medical students, both at the undergraduate and graduate levels, have minimal interest in learning about medical ethics (Mohamed et al., 2012). The dynamics of physician-patient communication seem to be a skill they gain over their career, which may explain why medical students are less satisfied with their relationship with patients than the more experienced physicians (Mohamed et al., 2012).

A survey of 805 attendees at the outpatient clinics at El-Demerdash University Hospital in Cairo showed that 81% of those attendees had limited comprehensive health literacy, and 84% had limited functional health literacy—the ability to understand and act on medical instructions (Almaleh et al., 2017). This means that many patients cannot digest the medical information even if it is shared with them. Moreover, the patient’s level of education might not allow the doctor to provide detailed information (Metwally et al., 2021).

## Methodology

### Sample

I selected 12 Egyptian doctor-influencers. All of them are popular physicians who are actively publishing health vlogs on YouTube. The sampling criteria were the following: 1) Language (vloggers had to create content in the Egyptian dialect of Arabic). 2) Content type (vloggers had to create content for social media users from the general public rather than for academic use). 3) Activity (vloggers had to be actively publishing on their channels/pages). I developed a list of 42 health professionals who fit these criteria by searching for health-related videos on YouTube using Arabic search terms (e.g., words such as “doctor” and the names of common medical specializations). I narrowed down the sample by filtering the channels and saving only those channels that met the inclusion criteria in a spreadsheet (e.g., by looking in the About section of their YouTube account or searching for other online profiles) and invited the health professionals to participate by sending invitation emails for virtual interviews.

Eventually, the sample included 12 Egyptian health professionals: two pharmacists, two psychologists, two pediatricians, a physiotherapist, a gynecologist, a gynecologist-obstetrician, and an orthopedic surgeon. Every participant had over 100,000 followers on their YouTube channel at the time of research; three of them had over 1 million followers.

### Semi-Structured Interviews

In April 2021, I conducted in-depth semi-structured interviews with the 12 health professional vloggers. All communications with participants were in Arabic, which optimized the clarity. The questions were asked in the Egyptian dialect, which the interviewer speaks fluently. This allowed the participants express themselves freely in their first language. All interviews were held virtually, via Zoom and WhatsApp. The interviews were used to examine the doctors’ self-described motivations and strategies for creating content and communicating with their audiences. I asked the doctor-vloggers direct questions about their communication of health research in vlogging, such as the following: Do you refer your audience to health research? Why/Why not? Do you leave a list of references for your audience on social media, where they can find the sources of your information? And how do you bring evidence to your content? The interview transcripts were organized by grouping the participants’ answers under three themes: 1) The significance of evidence (is it needed?) 2) Sharing evidence (when do you share it with the audience? Why do you or why do you not share evidence with the audience?) 3) Evidence reception (how are audiences using the evidence?)

***Critical Discourse Analysis (CDA)*** is a form of inductive analysis in which discursive events are instances of sociocultural practice, which are reflected on the levels of text, speech, and visuals (Fairclough, 1992). I sampled 48 videos from the YouTube channels and Facebook pages of the 12 participant doctor vloggers, using four videos per participant. The sample consisted of their most popular video, their most engaging video, and two other randomly selected videos. The videos are published on the participants’ YouTube and Facebook accounts, so there was no need for consent to analyze these public videos. I used the YouTube analysis tool developed by Influenex to assess the engagement rate of the videos from each channel. I used these metrics to find the most engaging videos. For the most popular videos, I used the YouTube API “Sort by most popular” to view the most popular videos from each channel. To facilitate coding, a questionnaire was created for organizing notes. Given the small sample (only 48 videos), the questionnaire was designed not for statistical analysis but for identifying the similarities and differences between the participants’ vlogs in terms of textual discourse (the use of the word “research”; the citation of medical research and the use jargons).

### Focus Groups

In September 2021, I convened two online focus groups titled Can Doctor Vloggers Mediate Health Research to Social Media Users? All 12 doctors were invited to participate in either one of these events based on their time preference. Instead of gathering all the doctor-influencers in one plenary focus group, I created mini focus groups of four or five participants to ensure that each doctor had a chance to share their commentary and insights. A mini focus group is a form of the standard focus group with a smaller number of participants (Krueger, 2014). It was hard to set a date and time that would be convenient for all the doctors, as some preferred a weekday evening, and others could only attend on the weekend, thus, one focus group was convened in the afternoon of a weekday and the other was organized in the evening of a weekend.

Each focus group lasted 90 minutes and was composed of two parts. In the first hour, I informed participants about the procedures of the study and presented findings from the interview analysis based on CDA. The participants were asked three questions: 1) What do you think about these results? 2) What do you agree/disagree with? 3) In your experience, what explains these results? For the remaining 30 minutes of each session, participants were engaged in a broader discussion about whether and how physician vloggers can mediate health research to social media audiences.

## Results

All the doctor-influencers agreed with the necessity of backing their content with evidence. Nevertheless, few of the 48 health vlogs created by the doctor-influencers included any reference to research. 40 vlogs did not mention the word “research” at all. The ones that did, generally focused on the research findings without stating the research methods or the year of publication.

In five videos, the doctor-influencer said “medical research suggests” without naming any specific studies. Only in two videos did the doctor-influencers mention the methodology and year of publication of the research they cited.

It was found during the interviews that the sources of evidence used in the vlogs differed across topics. Research becomes the main source of evidence for doctor-influencers when they vlog about new treatments, scientific discoveries, misinformation, and health myths. On the other hand, in the videos on basic health information, the doctors’ education and training, professional experience, and personal observations become the main sources of evidence. The number of years of work experience also played a role in the frequency of consulting research. There were two early-career doctor-influencers in the sample, both under 30 years old, and neither of them referred to their professional experience as evidence. Both participants stressed that they “study” the topics of their vlogs—a process that includes watching vlogs made by experienced physicians around the world, reading medical content about the topic, identifying controversial points, and consulting the latest research to finally form an opinion to pass on to the audience. Sometimes, these young doctor-influencers did not feel comfortable backing a certain medical argument. One of them explains:

> *I recognize my level of experience, so I am not yet at the level to support a study or refute it; I am slowly becoming a researcher. Maybe in a decade I will be able to decide my position in controversies; in the meantime, I prefer to explain them to my audience with my input, stressing that all the arguments are evidence-based*.

### 1. How Do Doctor-Influencers Present Evidence to Their Audiences on Social Media?

I found that the doctor-influencers used two criteria to decide what type of evidence to present in their vlogs: the topic of the vlog (as explained above) and the target audience.

#### Target Audience

Based on the demographic information they get from the social media platforms, and the comments they receive, the doctor-influencers believe they have 3 different types of audiences:

- *The Social Media Public:* The doctor-influencers believe this is the largest subset of the doctor-influencers’ audience. Also, the physicians believe that this subset of the audience wants the vlogs to be simple, informative, and a bit entertaining. Thus, they create health vlogs that simplify the medical content, inform, and entertain.
- *Fellow Physicians:* These are experienced physicians, pharmacists, and psychologists who may assess the participants’ vlogs in terms of the accuracy of and support for the health information they present. Therefore, in some vlogs the doctor-influencers cite health research as evidence to support their argument in front of their peers.
- *Early-Career Physicians:* According to the participants, early-career physicians form the smallest slice of the audience. They are young physicians looking to gain experience by watching medical vlogs. Therefore, the participants have made some videos describing medical examinations or procedures and the misdiagnoses of some illnesses. In the videos that target this subset of the audience, the physicians attach a bibliography to their videos to help the audience learn more about the topic.

While the first type of audience is perceived as mostly uninterested in checking health research and incapable of understanding it, this is not true of the other two groups. In fact, the participants’ awareness that their fellow doctors, pharmacists, and psychologists are watching their vlogs makes them keen to present evidence-based content.

#### The Topic of the Vlog

In addition to audience considerations, it was found during the interviews that the sources of evidence used in the vlogs differed across topics. The research was the main source of evidence for doctor-influencers when they vlogged about new treatments, scientific discoveries, misinformation, and health myths. On the other hand, in the videos introducing basic health information, the doctors’ studies, professional experience, and personal observations became their main sources of evidence. Specifically, thematic coding of the interviews and focus groups transcripts showed that the doctor-influencers become keen on citing health research in their vlogs and sharing a bibliography for the sources they used in the following cases: 1) Announcement of a major discovery in medicine or of a new treatment; 2) Correction of misinformation or health myths; 3) Self-defense against other vloggers (including doctors) who disagree with the vlogger; 4) The vlog content is or is expected to be controversial; 5) The vlog subject not the main area of expertise of the doctor.

#### Why Not Cite Research to the Audience?

##### It is “Not Even Wanted!”

Nevertheless, it was uncommon for the participants’ vlogs to cite academic research in the vlogs. Usually, they did not even list a bibliography in the caption box, because, according to the doctor-influencers, “the audience will not check the references” and references for medical research are “not even wanted.” In the focus groups, there was general agreement that the health vlogs’ viewers are very unlikely to check the references of a video by a doctor-influencer. The doctors justified this by saying, “I have not received a single comment requesting the bibliography so far” and

> *I usually check the number of clicks on the cards I put in my vlogs [cards are the on-screen hyperlinks for related videos created by the vlogger]; there is always just a small number of clicks on them; imagine what the number would be for external references.*

According to the doctor-influencers, this lack of interest in checking the references could be explained by the viewers’ inability to read academic papers; according to one participating psychologist, “we lack a research culture in the Arab world.” Furthermore, many among the general public audience have a low to medium level of education, and also some may have no education at all. One participant described a slice of their audience in an interview by saying,

> *Just expect that they [the social media audience] are good at reading and writing. I know for a fact that not all the audience are illiterate, With this level of knowledge, should I cite research?*

Another barrier for the audience to checking medical references is the language, as English is the dominant publication language in medicine, a foreign language in Egypt, which is an Arabic-speaking country. Plus, as some doctor-influencers noted, the medical journals are not easy to read for many people:

> *Even if a well-educated person with good English language skills went to a medical journal to double-check the information, they would not understand most of the content. The terms and illustrations there are not accessible for the public; only physicians and medical students will get them.*

The age of the audience of some health vlogs was an additional reason for the lack of research citations in the doctor-influencers’ vlogs, as a young audience would not be able to understand them. A psychologist-influencer explained in the focus group:

> *I am addressing people from different age groups, including teenagers; they definitely do not want to see “evidence,” and even if it is presented to them, they will not be able to understand it.*

Additionally, full descriptions of health research may leave the audience confused, and affect the doctor’s credibility, as one physician explained:

> *We [doctor-influencers] should not tell the audience about medical controversies, because they will get lost, and most probably will think the doctor indecisive.*

##### It is “Not Always Good!”

According to the participants, encouraging the social media users to visit medical research publications or even going “too far” in explaining medical facts to social media audiences might lead to the following:

- Incorrect self-diagnosis: the audience may start to link their minor health problems to symptoms of a serious disease (this is known as a somatic symptoms disorder).
- Incorrect diagnosis of others.: learning more about the anatomy of the human body, illnesses, and treatment could leave the audience feeling better informed than their peers who lack this knowledge. Therefore, the viewers would feel they are in a position to help others instead of going to a real doctor.
- Hypochondria, or the fear of having a physical illness, especially in cases when the physician describes common non-communicable illnesses (which happen without infection) or asymptomatic diseases (which happen without noticeable symptoms).
- Seeing the complications of a disease in its last stages, especially when pictures of the worst cases of the disease are available on the internet. A person who is newly infected with the disease may read medical sources about it and learn about these complications. The person may incorrectly think that they are close to this late stage of the disease, and may therefore stop seeking treatment. A psychologist gave an example of one of her patients who was diagnosed with paranoia: “she read in Wikipedia that a paranoia patient never fully recovers, which has affected her psyche and her faith in treatment.”

### 2. If Evidence is Not Research, What Is It?

The doctor-influencers stressed that they are always sharing evidence-based health information with their audiences and mediating verified medical content, but the form evidence that they are sharing is not always explicit. The evidence they bring to their audiences is not research, but the following:

#### a. Verified medical content

A participant explained in the focus group:

> *None of us [physicians] would risk our professional reputation by making inaccurate content. I believe all the reputable participants are, like me, verifying the information they use in their vlogs, even though they are sharing it with the social media audiences.*

#### b. Professional identity

The professional identity of the doctor-influencer is what makes the audience assume that the information published in a vlog is backed by research. In an interview, one doctor-influencer emphasized the link between the profession and evidence:

> *Should we really be worried about the audience not double-checking what doctors say in their vlogs? I do not think we should; are we concerned about this when this audience comes to us as patients and seeks help? Should we encourage them to verify our medical advice before following it?*

Therefore, the doctor-influencers tend to emphasize their professional identity by, for instance, dressing professionally: in 33 of the videos, the physician was wearing a white coat or a hospital gown, highlighting their credibility and emphasizing their identity as medical professionals. Many of those who did not appear in a coat or gown highlighted their professionalism by wearing a suit or formal blazer. The physicians’ credibility in their profession is the evidence they present to their social media audience.

In fact, most of the doctor-influencers in this research, particularly the mid-career and established physicians, did not see a robust link between citing research to their audiences and maintaining or increasing their credibility. In one of the focus groups, when I raised a question about whether the lack of research citations affected the doctor-influencer’s credibility, one gynecologist said, “I am credible! I do not really need to prove it to the audience.” This participant is a professor at the Cairo Faculty of Medicine and has a good reputation in his specialization; this is his source of credibility, which precedes vlogging.

The doctor-influencers used different strategies to prove their credibility to audiences. One of them was listing their degrees and professional credentials. Every participant shared a full list of their degrees, training certificates, and honors in the About section of their YouTube channel and Facebook page to emphasize their credibility as a physician.

There were also some physicians who believed that credibility is all about good communication of content, saying,

> *Credibility is not built by citing research; do you think patients nowadays recommend doctors based on the number of certificates on the walls of their clinics? No, they trust the doctor if his workplace looks fancy and he is a good communicator.*

A growing audience was taken as an indicator of credibility: “People would not have subscribed to my [YouTube] channel and watched my vlogs if they doubted my credibility”; “I get patients in my practice who come after watching my vlogs; they would not have done that if I had not won their trust.”

Lastly, the doctor-influencers counted on the social media audience to search for evidence when they wanted it, which they would likely do using research engines. As one doctor-influencer explained,

> *If the social media users were curious to know more about what I am saying, or to double-check it, they would just research the topic on Google or Wikipedia, not in research papers.*

Another way would be triangulating the information in one vlog with others made by different physicians, as one participant explained:

> *It is not only me who is vlogging; there are many physician-vloggers; if a viewer wanted evidence, I think they would watch vlogs by different doctors and see what they agree on.*

## Discussion

The doctor-influencers agreed on the necessity of backing their content with evidence. They stressed in the focus groups that they are always sharing evidence-based health information with their audiences by mediating verified medical content, but the form of evidence that they are sharing is not always explicit. Although the doctors said that they read and consider evidence when planning and creating their content, they rarely presented that evidence to the audience. The doctor-influencers believed that their professional identity automatically makes the audience assume that the information in their vlogs is backed by research.

We cannot isolate the influence of dominant cultural practices from the findings. The participants’ justifications for not relying on research as evidence matches the circumstances of high illiteracy and low health literacy, which makes no point in referring to scholarship because it will not be clear to the majority of the audience. These findings highlight the challenges of science communication in societies with a low level of education. In these settings, communicators may need to move beyond mediating research studies on social media and instead focus on simplifying, translating, and connecting evidence-based information to the cultural context.

The implications of the reliance on the professional identity rather than research as evidence of the accuracy of the health information may lead to unethical practices. For instance, people could take advantage of it to spread misinformation, or some physicians to make content outside of their specializations for commercial purposes. In this case, the audience is very likely to believe the physician and accept the information, advice, or recommendations from them without verification simply because they are “a doctor.” More research is needed on the downsides of sharing health research with the public through social media, especially in the absence of affordable health care services, where people prefer self-diagnosis and peer consultation to visiting a physician.

## Acknowledgments

I would like to acknowledge Dr. Juan Pablo Aplerin, Alice Fleerackers, and Dr. Lauren Maggio for their support in scoping this paper.

## References

Adebayo, C. T. (2021). Physician-patient interactions in Nigeria: A critical-cultural perspective on the role of power. Journal of Intercultural Communication Research, 50(1), 21–40.

Almaleh, R., Helmy, Y., Farhat, E., Hasan, H., & Abdelhafez, A. (2017). Assessment of health literacy among outpatient clinics attendees at Ain Shams University Hospitals, Egypt: A cross-sectional study. Public Health, 151, 137–145.

World Bank, W. (2017). UNESCO Institute for Statistics. Arab Rep. https://data.worldbank.org/indicator/SE.ADT.LITR.ZS?locations=EG

World Bank, W., & World Health Organization. (2017). Hospital Beds per 1,000 people. Arab Rep.

Emanuel, E. J., & Emanuel, L. L. (1992). Four models of the physician-patient relationship. JAMA, 267(16), 2221–2226.

Fairclough, N. (1992). Discourse and text: Linguistic and intertextual analysis within discourse analysis. Discourse & Society, 3(2), 193–217.

Hassig, K. (2021). Cultural health capital and the doctor-patient encounter: An exploratory analysis. [published thesis]. University of Oklahoma

Jiang, S., & Liu, J. (2020). Examining the relationship between Internet health information seeking and patient-centered communication in China: Taking into account self-efficacy in medical decision-making. Chinese Journal of Communication, 13(4), 407–424.

Kabbash, I., Rania, E. S., Zayed, H., Alkhyate, I., Omar, A., & Abdo, S. (2021). The brain drain: Why medical students and young physicians want to leave Egypt. Eastern Mediterranean Health Journal, 27(11), 1102–1108.

Krueger, R. A. (2014). Focus groups: A practical guide for applied research. Sage publications.

Langford, A. T., Roberts, T., Gupta, J., Orellana, K. T., & Loeb, S. (2020). Impact of the internet on patient-physician communication. European Urology Focus, 6(3), 440–444.

Metwally, A. M., Amer, H. A., Salama, H. I., Abd El Hady, S. I., Alam, R. R., Aboulghate, A., & Eldali, A. (2021). Egyptian patients’/guardians’ experiences and perception about clinical informed consent and its purpose: Cross sectional study. Plos One, 16(6), 0252996.

Mohamed, A. M., Ghanem, M. A., & Kassem, A. A. (2012). Knowledge, perceptions and practices towards medical ethics among physician residents of University of Alexandria hospitals, Egypt/Connaissances, perceptions et pratiques en matiere d’ethique medicale des internes des centres hospitaliers universitaires d’Alexandrie (Egypte). Eastern Mediterranean Health Journal, 18(9), 935.

Riahi, H., Bazmi, S., Enjoo, S. A., Ahmadnia, S., & Afshar, L. (2020). The physician-patient relationship: How it is represented in Iranian national television. Journal of Medical Education, 19(1), 2107905.

Shawky, S. (2018). Measuring geographic and wealth inequalities in health distribution as tools for identifying priority health inequalities and the underprivileged populations. Global Advances in Health and Medicine, 7, 2164956118791955.

Shalaby, S., Saied, M, & Zaid, H. (2019). Physician-patient communication: Perception and practice among doctors working in Tanta University outpatient clinics, Egypt. Egyptian Journal of Occupational Medicine, 43(3), 453–467.

Suarez-Lledo, V., & Alvarez-Galvez, J. (2021). Prevalence of health misinformation on social media: Systematic review. Journal of Medical Internet Research, 23(1), 17187.

Sylvia Chou, W. Y., Gaysynsky, A., & Cappella, J. N. (2020). Where we go from here: Health misinformation on social media. American Journal of Public Health, 110(S3), 273–275.

Szasz, T. S., & Hollender, M. H. (1956). A contribution to the philosophy of medicine: The basic models of the doctor-patient relationship. AMA Archives of Internal Medicine, 97(5), 585–592.

Unger, J. P., Ghilbert, P., & Fisher, J. P. (2003). Doctor-patient communication in developing countries. BMJ, 327(7412), 450–450.

Huo, J., Desai, R., Hong, Y. R., Turner, K., Mainous III, A. G., & Bian, J. (2019). Use of social media in health communication: findings from the health information national trends survey 2013, 2014, and 2017. Cancer Control, 26(1), 1073274819841442.

Vraga, E. K., & Bode, L. (2020). Correction as a solution for health misinformation on social media. American Journal of Public Health, 110(S3), 278–280.

